# Cell profiling of mouse acute kidney injury reveals conserved cellular responses to injury

**DOI:** 10.1101/2020.03.22.002261

**Authors:** Yuhei Kirita, Haojia Wu, Kohei Uchimura, Parker C. Wilson, Benjamin D. Humphreys

## Abstract

After acute kidney injury (AKI), patients either recover or alternatively develop fibrosis and chronic kidney disease. Interactions between injured epithelia, stroma and inflammatory cells determine whether kidneys repair or undergo fibrosis, but the molecular events that drive these processes are poorly understood. Here, we use single nucleus RNA sequencing of a mouse model of AKI to characterize cell states during repair from acute injury. We identify a distinct proinflammatory and profibrotic proximal tubule cell state that fails to repair. Deconvolution of bulk RNA-seq datasets indicates that this “failed-repair proximal tubule cell” or FR-PTC, state can be detected in other models of kidney injury, increasing in the aging rat kidney and over time in human kidney allografts. We also describe dynamic intercellular communication networks and discern transcriptional pathways driving successful vs. failed repair. Our study provides a detailed description of cellular responses after injury and suggests that the FR-PTC state may represent a therapeutic target to improve repair.

**Significance Statement:** Single nucleus RNA sequencing revealed gene expression changes during repair after acute kidney injury. We describe a small population of proximal tubule cells that fail to repair (FR-PTC). Since this subpopulation expresses abundant pro-inflammatory and profibrotic genes, it may represent a new therapeutic target to improve repair and reduce fibrosis after AKI.

## Introduction

Kidneys maintain fluid and electrolyte balance, excrete waste products and regulate blood pressure. Composed of approximately one million functional units called nephrons, the kidneys receive ∼20% of cardiac output. Nephrons have high metabolic activity rendering them susceptible to injury from toxins or reduced blood flow. These insults cause acute kidney injury (AKI) characterized by decreased kidney function. In early stages of AKI, epithelial cells die and surviving epithelia dedifferentiate accompanied by inflammation. Dedifferentiated epithelial cells then proliferate and redifferentiate to repair the damaged nephron (1, 2). There are no specific treatments for acute kidney injury but with supportive care the kidney’s intrinsic repair capacity usually allows functional recovery over a period of days to weeks.

After repair, kidney function may not return back to baseline due to residual subclinical inflammation and fibrosis. Survivors of AKI are at high risk of developing future chronic kidney disease and even kidney failure (3). The mechanisms for failed repair are not well understood, but a sub-population of injured proximal tubule epithelia (the segment most susceptible to injury) are proposed to become arrested at the G2 / M cell cycle phase and adopt a senescence-associated secretory phenotype (4). This may prevent complete repair, driving inflammation and fibrosis, and mouse ischemia-reperfusion injury (IRI) models this process well (5). The aim of our study was to understand the cellular events underlying both recovery from AKI as well as the transition to chronic kidney disease. Bulk transcriptional profiling has successfully characterized kidney injury and recovery (5, 6), but these approaches describe a transcriptional average across cell populations, which may hide or skew signals of interest. We hypothesized that understanding transcriptional changes in single cell types over the course of AKI, repair and fibrosis would provide unique insights into disease pathogenesis and potentially identify new therapeutic strategies.

## Results

We performed single nucleus RNA-sequencing (snRNA-seq) on cryopreserved mouse kidney (7). Mice were sacrificed at 4 and 12 hours and 2, 14 and 42 days after bilateral ischemia reperfusion injury (IRI) (Fig. 1A). Both histologic changes (Fig. S1) and blood urea nitrogen (Fig. 1B) levels confirmed acute injury and its resolution in mouse.

**Figure 1.**
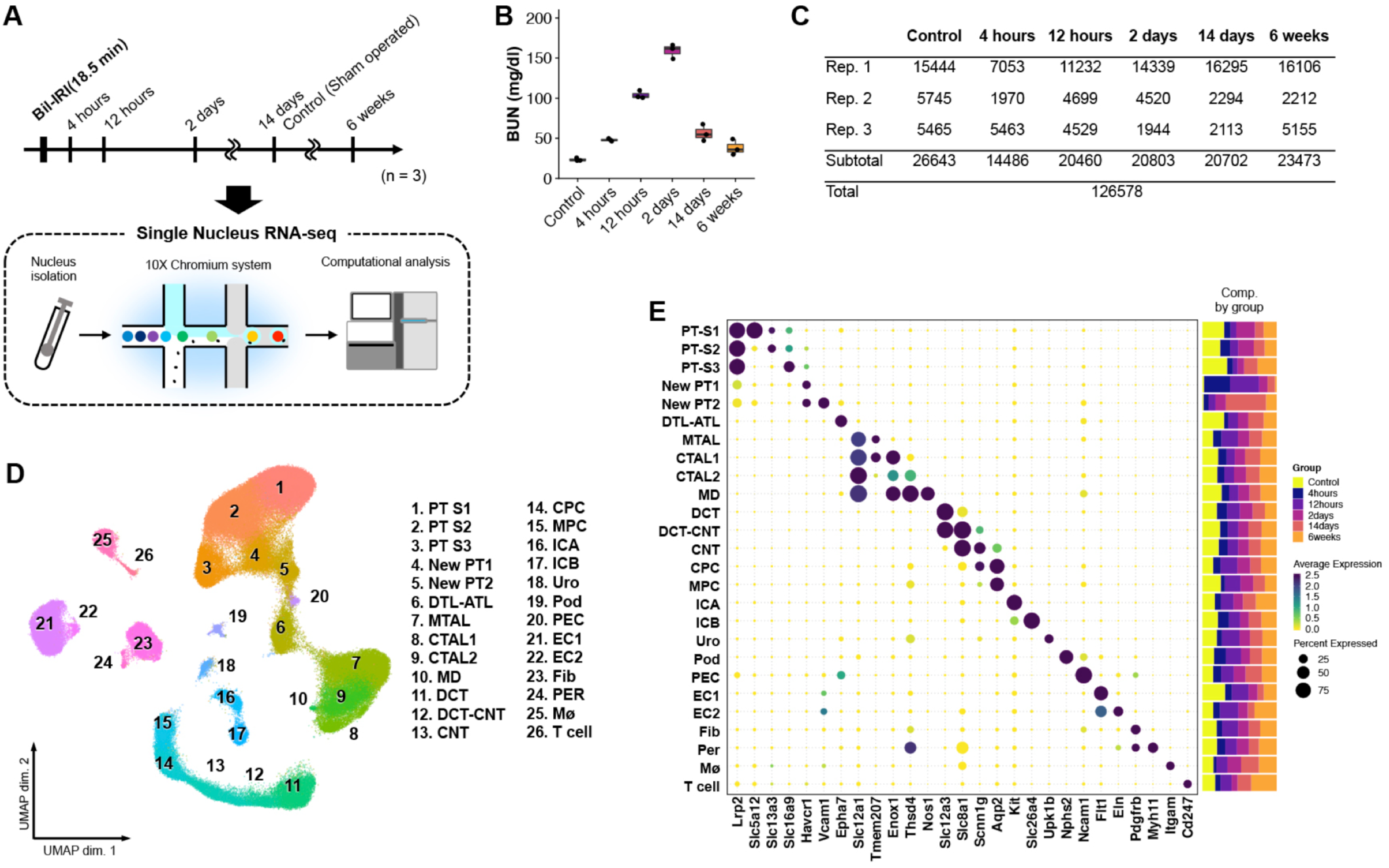
Single-nucleus RNA-seq atlas of mouse IRI kidney. (A) Summary of experimental strategy. n = 3 mice per group. (B) Blood urea nitrogen (mg/dl) after sham and IRI. Data are shown as the mean ± SEM. (C) Table with details of group, replicates, and cell numbers of mouse IRI datasets present in this figure. (D) Umap plots of all mouse IRI kidney datasets integrated with Harmony. PT-S1, S1 segment of proximal tubule; PT-S2, S2 segment of proximal tubule; PT-S3, S3 segment of proximal tubule; DTL, descending limb of loop of henle; ATL, thin ascending limb of loop of henle; MTAL thick ascending limb of loop of henle in medulla; CTAL thick ascending limb of loop of henle in cortex; MD, macula densa; DCT, distal convoluted tubule; CNT, connecting tubule; CPC; principle cells of collecting duct in cortex; MPC; principle cells of collecting duct in medulla; ICA, type A intercalated cells of collecting duct; ICB, type B intercalated cells of collecting duct; Uro, urothelium; Pod, podocytes; PEC, parietal epithelial cells; EC, endothelial cells; Fib fibroblasts; Per, pericytes; Mø, macrophages. (E) Dot plot displaying gene expression patterns of cluster-enriched markers, and bar plot displaying composition of clusters by groups.

After quality control filtering, we obtained 26,643 cells from healthy mouse kidneys. Visualization of single nucleus transcriptomes in UMAP space resolved 26 separate clusters (Fig. S2A, Dataset S1). Sub-clustering of both epithelial (descending loop of Henle, thin ascending limb) and non-epithelial cells (immune, endothelial, stromal) revealed additional cell clusters (Fig. S2B). For example, five separate endothelial clusters were identified including arterial, lymphatic, descending vasa recta and cortical vs. medullary endothelium. Eight stromal clusters were detected, including mesangium and *Ren1*-positive juxtaglomerular apparatus cells (Fig. S2B). Major cell types and subclusters were identified based on cell type-specific markers (Fig. S2C, D, Dataset S1) (8-10). Each nephron segment performs unique reabsorptive and secretory functions to transform filtrate into urine, and this is reflected by segment-specific expression of all detected solute linked carriers, ATPases and channels (Fig. S3).

### Proximal tubule responses to acute injury

We generated 99,935 mouse AKI single cell transcriptomes (Fig. 1C) and integrated these with the healthy datasets using the Harmony algorithm to reduce batch effects (Fig 1D) (11). We could define unique anchor genes for all clusters in the integrated datasets, and defined the relative abundance of each cluster in healthy vs. injured kidney. For example, the AKI kidneys contributed a much larger fraction of leukocytes than healthy kidneys (Fig. 1D, E). All original clusters were retained, but two new clusters in mouse (Fig. 1D cluster 4: New PT1 and cluster 5: New PT2) appeared in injury and these were located adjacent to healthy proximal tubule in UMAP space. Analysis of marker gene expression showed that these new clusters expressed the proximal tubule marker *Lrp2* encoding Megalin, but also the injury marker *Havcr1* encoding Kim1, indicating that these clusters represented an injured proximal tubule state in mouse.

We focused our analysis on proximal tubule, since this segment suffers the most injury due to high metabolic activity. Unsupervised subclustering of all mouse proximal tubule cells across timepoints yielded three healthy subclusters (the S1, S2 and S3 segments of the proximal tubule), one repairing subcluster, and three injured subclusters (Fig. 2A). Differential gene expression and gene set enrichment analysis (GSEA) were performed to characterize the subclusters (Dataset S2). Assigning timepoints to these clusters in UMAP space helps visualize temporal changes in proximal tubule gene expression (Fig. S4A).

**Figure 2.**
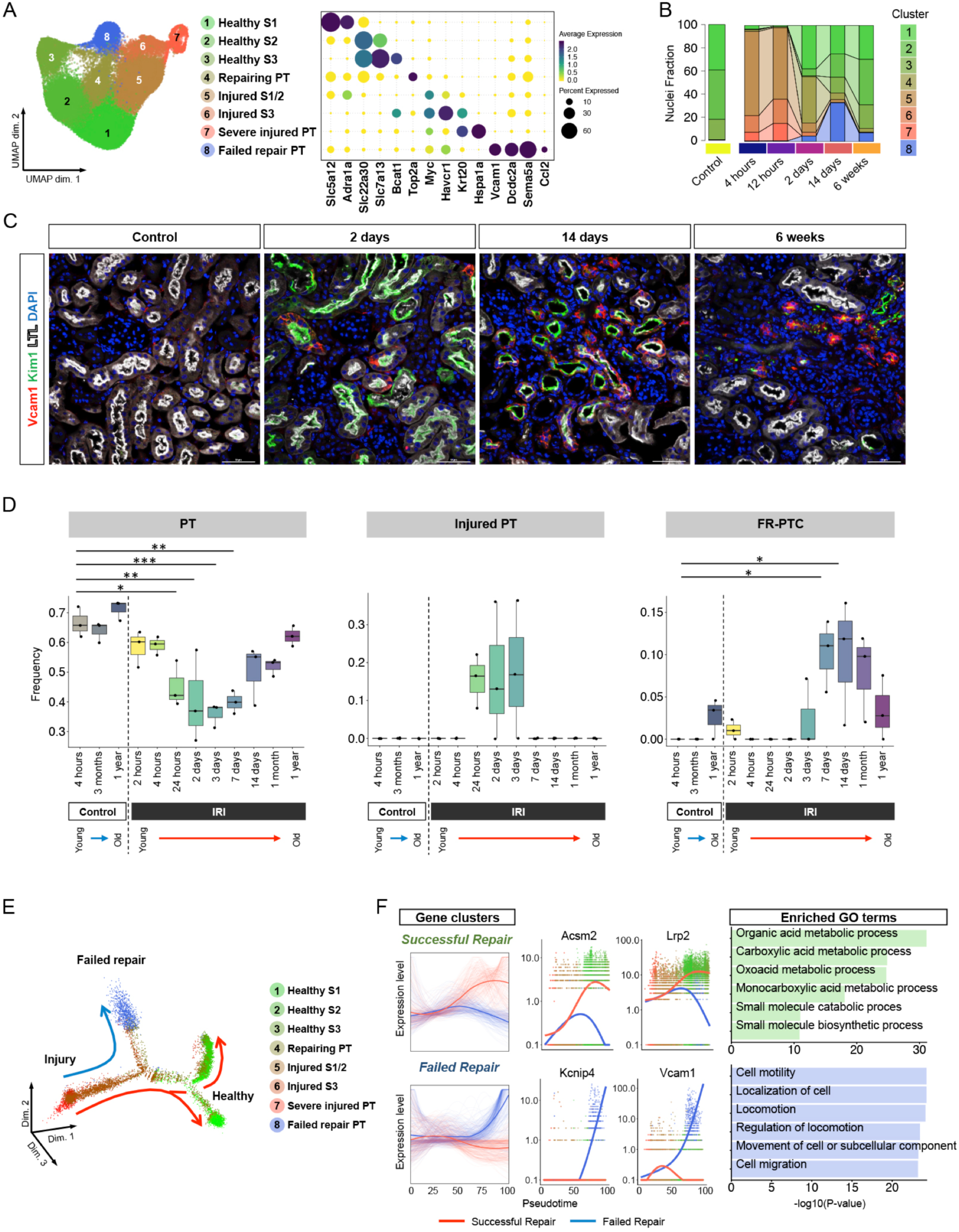
Time-course analysis of proximal tubular cells revealed new cell state, failed repair proximal tubular cells. (A) Umap displaying the clustering of proximal tubular cells without Harmony integration and dot plot displaying gene expression patterns of cluster-enriched markers. (B) Bar plot displaying composition of groups by clusters. (C) Representative images of immunofluorescence staining for VCAM1 (red), Kim1 (green) and LTL (white). (D) Deconvolution analysis of bulk RNA-seq mouse kidney IRI dataset using gene sets specific for healthy PT, injured PT and failed repair PT. *P < 0.05; **P < 0.01; ***P < 0.001, one-way ANOVA with post hoc Dunnett’s multiple comparisons test.(E) Monocle2 pseudotime trajectory of proximal tubular cells colored by cluster identity. (F) Gene expression dynamics on the trajectories. The expression dynamics of DEGs were cataloged into 3 clusters across pseudotime shown as red lines (successful repair) and blue lines (failed repair). Thick lines indicate the average gene expression patterns in each cluster. The top 6 enriched gene ontology terms for each cluster are shown on the right.

There were three categories of injured proximal tubule cells – we annotated these as ‘injured S1/2’, ‘injured S3’ and ‘severe injured PT’ respectively. ‘Injured S1/2’ and ‘injured S3’ were primarily composed of cells from the 4- and 12-hour timepoints, and expressed *Myc*, which encodes c-Myc playing a role in cell cycle progression, and *Havcr1*, and also shared a part of differentially expressed genes (DEGs) of ‘healthy S1’ and ‘healthy S2’ as its DEGs (Fig. 2A, Dataset S2) respectively. ‘Severe injured PT’ shared expression of many injured PT genes, but additionally expressed the tubule injury markers *Krt20* (5, 6), as well as genes encoding heat shock protein, suggesting a more severe injury to these cells. GSEA showed that these proximal tubule injury states had enrichment of response to stress and damage, and ‘severe injured PT’ had additionally had “cell cycle arrest” (Fig. S4B).

A ‘Repairing PT’ cluster arose two days after injury and had enrichment of “mitotic cell cycle” and “meiotic cell cycle” terms, including up-regulation of *Top2a* which is essential for proliferation. Cell cycle status analysis revealed that ‘repairing PT’ had the highest proportion of cycling cells (Fig.S4C, D). In contrast, the proximal tubule injury clusters had almost disappeared by two days (Fig. 2B), and a new distinct cell cluster arose, growing and reaching nearly 30% of all proximal tubule states at 14 days after injury, and remaining ∼8% of total proximal tubule by 6 weeks after injury (Fig. 2B). This cluster expressed a distinct set of genes not observed in either healthy or acutely injured mouse proximal tubule. These included *Vcam1, Dcdc2a* and *Sema5a* (Fig. 2A, Dataset S2). Because this cluster additionally downregulated expression of terminal differentiation markers such as *Slc5a12, Slc22a30* and *Slc7a13* even at late timepoints, we annotated this cluster as ‘failed repair proximal tubule cells,’ or FR-PTC. We recently reported that ∼20% of injured proximal tubule cells fail to repair at two weeks after AKI, and the presence of the FR-PTC cluster in the current analysis supports and extends those results (12). GSEA of FR-PTC revealed terms such as “positive regulation of lymphocyte activation”, “NIK NFĸB signaling” and “cell cell signaling by Wnt”, suggesting FR-PTC are pro-inflammatory (Fig. S4B). We localized FR-PTCs after IRI by immunofluorescence. Vcam1 positive FR-PTCs emerged within Kim1 positive injured tubule in a scattered manner at 2 days after IRI, then expanded and remained within atrophic tubules with or without Kim1 expression at late timepoints (Fig. 2C).

Mapping the PT subclusters back onto the entire dataset revealed that the “New PT1” cluster was primarily composed of the three acute injury states and the “New PT2” cluster was primarily composed of FR-PTC (Fig. S4E, F). We therefore annotatedthese clusters as “Injured PT” and “FR-PTC” respectively in further analyses. We asked whether these distinct PT states could be detected in other bulk RNA-seq datasets by applying the BSEQ-sc deconvolution algorithm (13). We assessed the fraction of healthy PT, injured PT and FR-PTC in an independent mouse IRI one year time course (5). This showed that healthy PT decreased after injury, but gradually recovered with time, injured PT increased after injury but resolved by 7 days and FR-PTC appeared beginning one week after injury and persisted (Fig. 2D). Similar trends were observed in a mouse folic acid injury model (Fig. S5A)(14). In human protocol biopsies from kidney transplants, FR-PTC increased at one year after transplant compared to pre-transplant (Fig. S5B) (15). The proportion of FR-PTC also increased with age in rat kidneys, increasing from ∼5% at six months to ∼12% at 27 months (Fig. S5C) (16).

We reconstructed proximal tubule lineage relationships during repair by pseudotemporal ordering. The mouse trajectory began with injury, and most cells progressed to healthy S1/S2 or S2/S3 proximal tubule segments, but FR-PTCs formed an alternate branchpoint off the successful repair trajectory, indicating that FR-PTCs represent a distinct cell state (Fig. 2E). Gene ontology analysis across the pseudotime trajectory showed that the successful repair trajectory included terms that would be expected in cells that are re-differentiating such as “organic acid metabolic process” and “carboxylic acid metabolic process” (Fig. 2F). The FR-PTC arm included terms like “cell motility” and “cell migration.” These results define FR-PTC as a distinct state after injury characterized by a unique set of markers and that persists after resolution of injury.

We next used single-cell regulatory network inference and clustering (SCENIC) to map the gene regulatory networks governing these proximal tubule cell states (17). We discovered marked differences in regulon activity between FR-PTC, and either healthy or acutely injured states, providing further evidence that these are distinct proximal tubule cell states (Fig. 3A, Dataset S3). The FR-PTC cluster had regulon activity for both Relb and NFkB, suggesting a proinflammatory status for these cells (Fig. 3B). Also specific to the FR-PTC cluster was the Tcf7l1 regulon, which mediates Wnt signaling, consistent with the strong Wnt GSEA terms in this cluster. Proximal tubule canonical Wnt signaling is important both in specification and development, but also in disease (18, 19).

**Figure 3.**
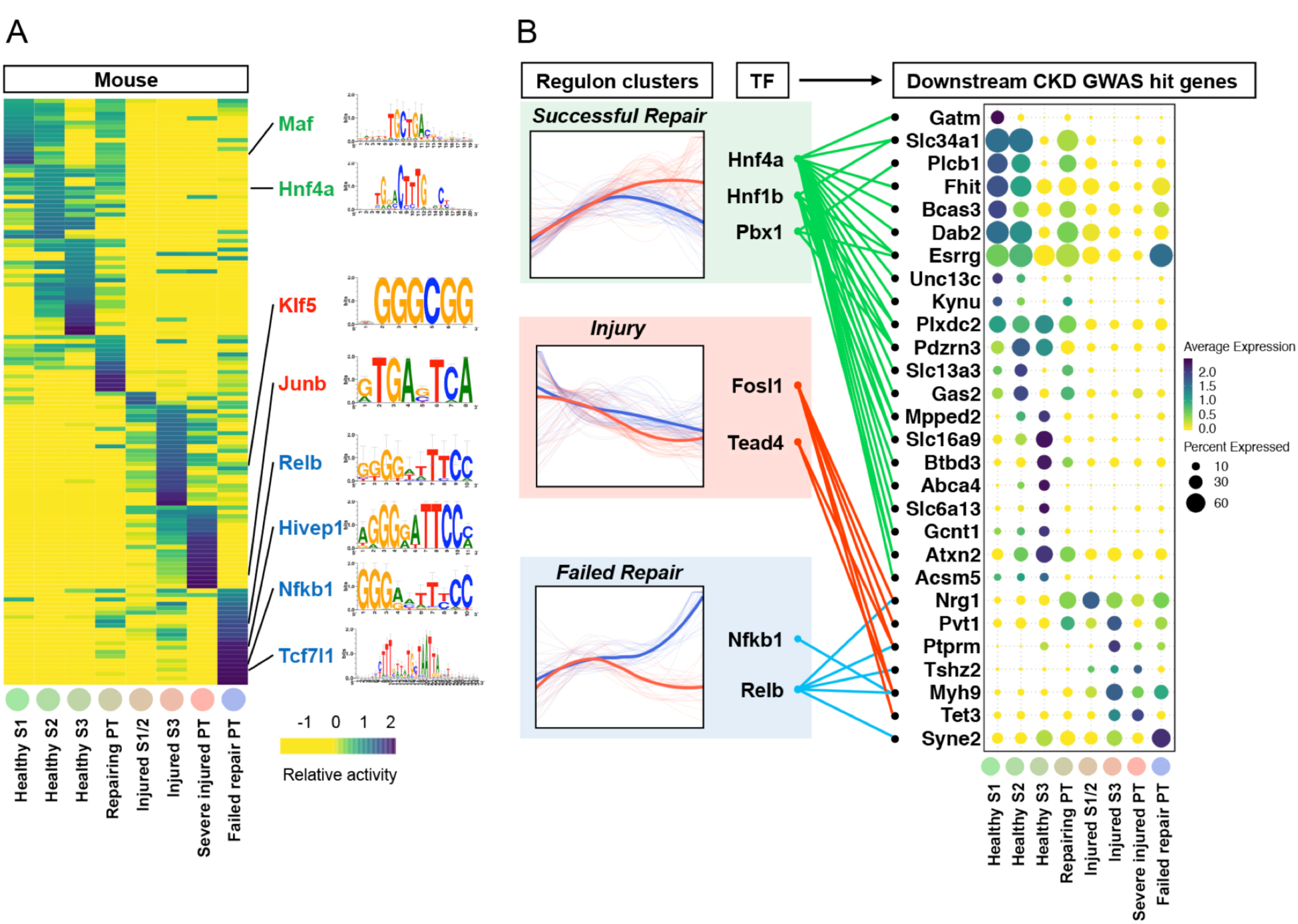
Gene regulatory network analysis of proximal tubular cells predicts transcription factors for successful repair and failing repair. (A) Heat map depicting the average regulon activity in each cluster of proximal tubular cells. Representative transcription factors are highlightened along with corresponding DNA-binding motifs. Cluster identities are according to Fig.2. (B) Regulon activity dynamics on the pseudotime trajectory. The regulon activity dynamics were cataloged into 3 clusters across pseudotime shown as blue lines (successful repair) and red lines (delayed repair). Thick lines indicate the average gene expression patterns in each cluster. Pseudotime trajectories are according to Fig.2. Transcription factors that have downstream CKD GWAS hit genes are displayed with their target genes expression patterns by dot plots.

We mapped the relationship between transcription factors identified by this analysis and their regulation of GWAS genes associated with CKD. In the successful repair cluster, *Hnf4a, Hnf1b* and *Pbx1* drive expression of multiple differentiation-associated genes that are also GWAS hits for CKD including a variety of solute linked carriers plus *Plxdc2, Gas2* and *Dab2* (Fig. 3B) (20). By contrast, both injury and the FR-PTC clusters had strong gene regulatory network signals for transcription factors regulating the expression of GWAS genes that were not expressed in healthy proximal tubule but rather in the injured state. Examples include the non-muscle myosin gene *Myh9*, present in both injured and delayed repair clusters, and *Nrg1* encoding the epidermal growth factor ligand neuregulin, present primarily in the injured clusters. In particular, NFkB and Relb regulons were specific to FR-PTC, and we could map downstream CKD GWAS genes to specific clusters, both healthy and injured. These results provide functional annotations of cell state-specific transcription factor mediated regulatory networks, helping to elucidate the cellular context for susceptibility loci identified in CKD GWAS studies.

We could detect FR-PTC marker expression in apparently healthy human kidneys. Consistent with a prior report, these cells are located in a ‘scattered’ fashion, adjacent to normal proximal tubule cells, throughout the proximal tubule (21). Examination of images from the Human Protein Atlas (22) shows scattered cells in healthy human kidney that express VCAM1 and DCDC2 (Fig. S6A). We could also detect evidence for downregulation of differentiation markers in isolated cells scattered throughout the nephron as well (Fig. S6B). These results suggest that a conserved injury response occurs in individual, isolated cells even during homeostasis.

### Stromal cell responses to injury

Recent scRNA-seq analyses have revealed unexpected stromal heterogeneity in both developing and adult kidney (8, 9, 23). With Harmony integration, we combined the stromal clusters from all timepoints to identify eight stromal cell subclusters (Fig. 4A). These included four fibroblast populations that differed according to their cortical or papillary site of origin. We could also detect a pericyte and vascular smooth muscle cell population that were both characterized by strong expression of Notch pathway constituents such as *Notch3* and *Jag1*, consistent with important roles for this pathway in pericyte development and angiogenesis (24, 25). We identified renin-secreting juxtaglomerular cells as well as mesangial cells. Several stromal clusters differed according to kidney region. For example, the cortical fibroblast marker *Dapk2* was expressed in cluster 3 and 4 but not in cluster 1 and 2 (Fig. 4A) (26), suggesting that cluster 3 and 4 are cortical fibroblasts and cluster 1 and 2 are medullary fibroblasts.

**Figure 4.**
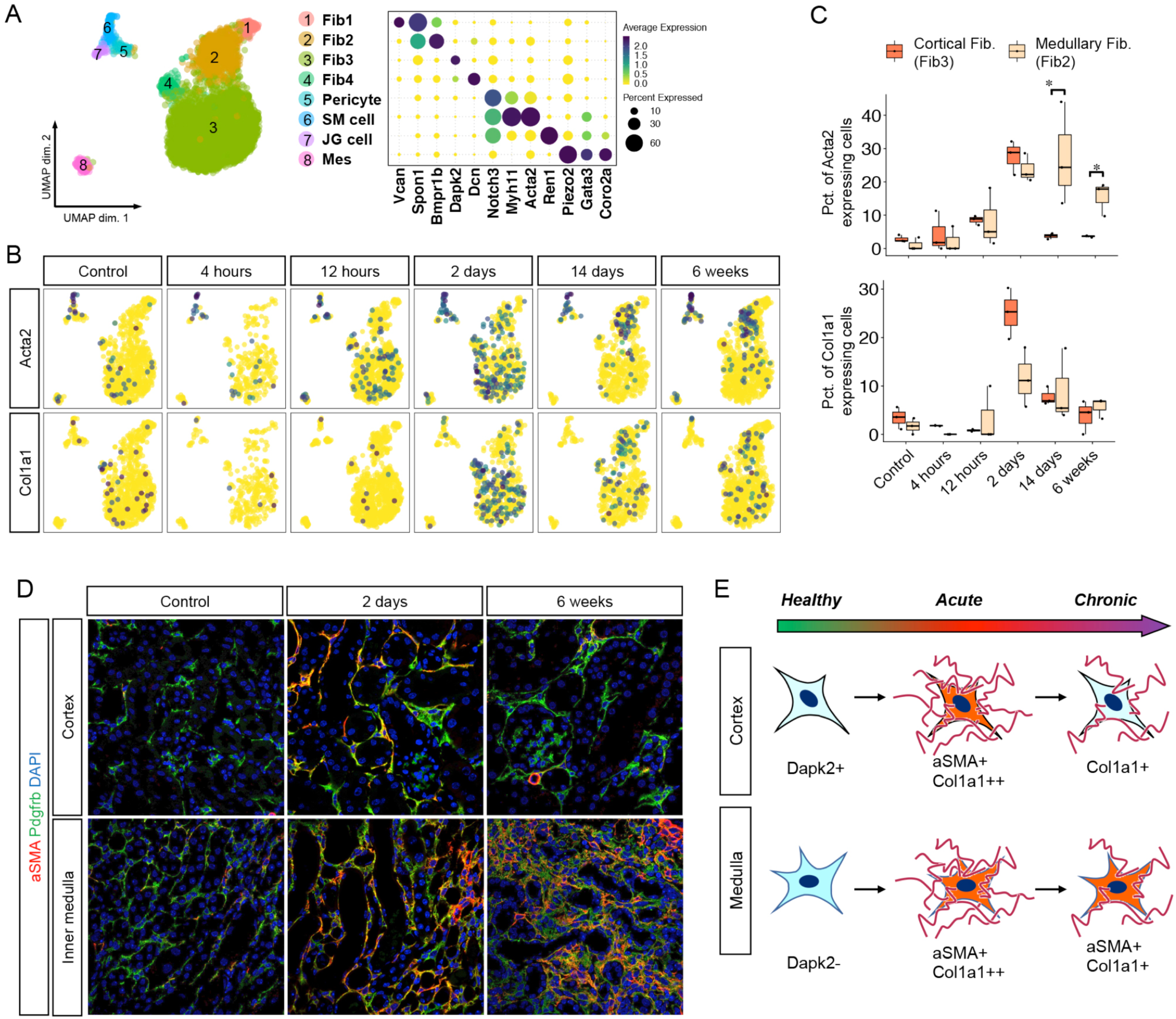
Novel stromal subtypes are identified including a population with reversible expression of αSMA. (A) Umap displaying the clustering of all stromal cells with Harmony integration and dot plot displaying gene expression patterns of cluster-enriched markers. (B) Umap displaying expression levels of typical myofibroblast markers in each group of the datasets. (C) Boxplot displaying percentages of myofibroblast markers expressing cells in mouse fibroblast3 (cortical fibroblast) and fibroblast2 (medullary fibroblast) at each time point. (D) Representative images of immunofluorescence staining for aSMA (red) and Pdgfrb (green). (E) Diagram of the fate of fibroblasts after IRI.

Myofibroblasts secrete matrix proteins and are critical for fibrogenesis (27). Two marker genes for myofibroblasts are *Acta2* and *Col1a1*. In healthy kidney, *Acta2* expression was largely restricted to smooth muscle cells. After IRI, there was strong upregulation of *Acta2* across all stromal clusters with the exception of mesangial cells, and *Col1a1* was also strongly induced in fibroblasts (Fig. 4B). We could observe that cortical fibroblasts only transiently upregulated *Acta2* and *Col1a1* with a peak at day 2 after IRI, whereas medullary fibroblasts showed sustained expression of *Acta2* but not *Col1a1* at 6 weeks (Fig. 4B, C). Medullary fibroblasts also increased as a fraction of the total stromal cells over time (Fig. S7). These results suggest an unappreciated plasticity of kidney stroma. We could verify the injury-induced transient upregulation of a-smooth muscle actin (αSMA), the protein encoded by *Acta2*, in cortical fibroblasts but not medullary fibroblasts, by immunofluorescence analysis (Fig 4D). These results suggest regional differences in the response of fibroblasts to injury, with medullary fibroblasts progressing to a myofibroblast cell state and cortical fibroblasts reverting to their prior quiescent state (Fig. 4E).

### Ligand-receptor interactions during injury and repair

Finally, we leveraged our datasets to explore how injury affects intercellular communication within the kidney. We performed ligand-receptor analysis across all time points with simplified global clustering (Fig. 6A). We highlight the tubulointerstitial compartment, comprising proximal tubule, endothelium, stroma and leukocytes, because interstitial fibrosis in the kidney cortex (where proximal tubule is located) best predicts future kidney failure (28). *Ccl2* and its receptor *Ccr2* play important roles in AKI by recruiting monocytes and T cells (29). We used a standardized ligand-receptor score to quantitate signaling from *Ccl2* in tubulointerstitium to *Ccr2* in leukocytes across time (30). This revealed a temporal progression whereby fibroblasts and endothelial cells were the first cell type to signal to leukocytes, followed by leukocyte-leukocyte signaling at day 2, and finally increasing Ccl2 – Ccr2 signaling from FR-PTC (Fig. 5B, C). Compared to FR-PTC, proximal tubule destined for successful repair minimally upregulated Ccl2 even in acute injury, emphasizing the pro-inflammatory nature of FR-PTC.

**Figure 5.**
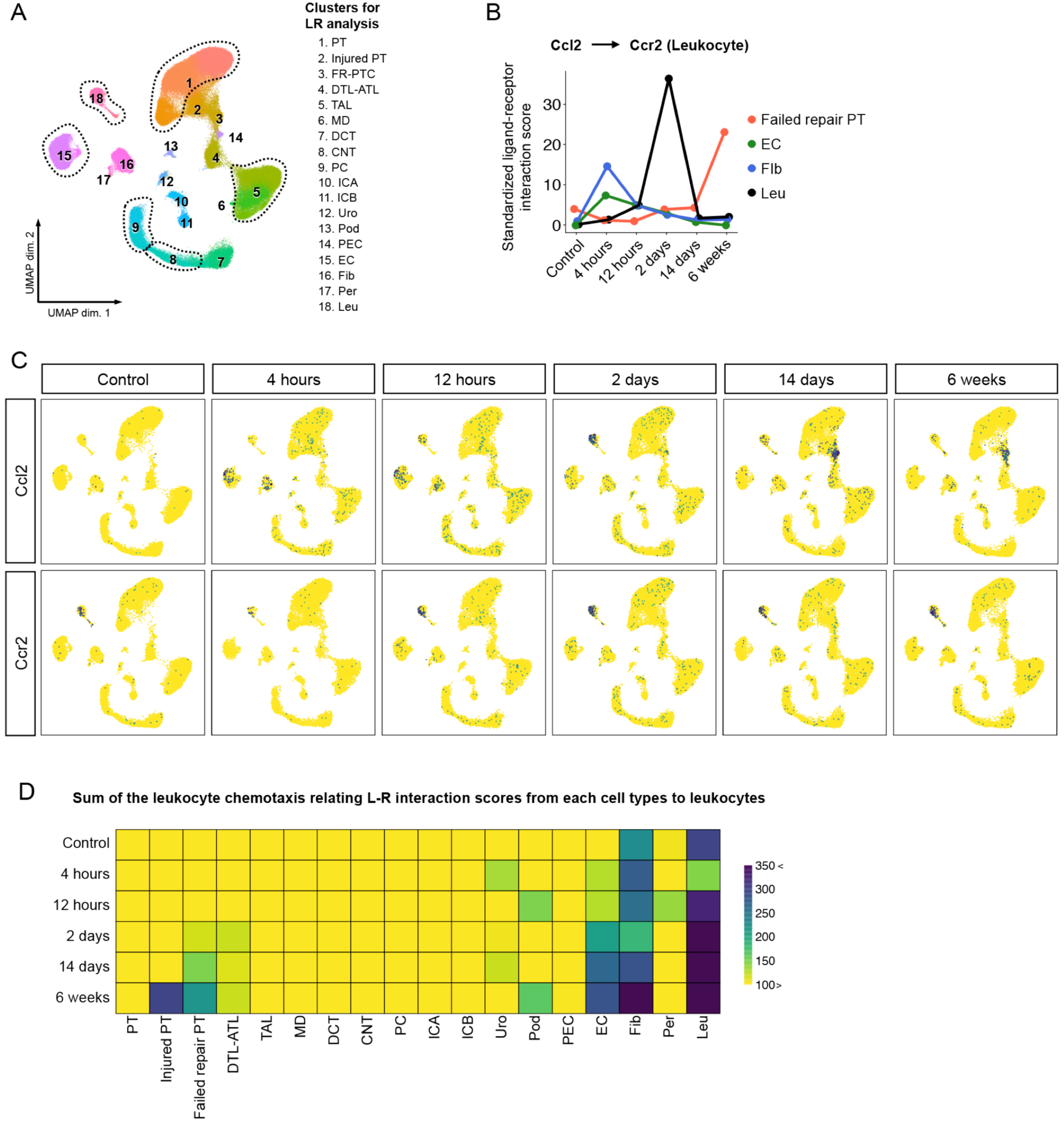
Ligand-receptor analysis reveals dynamics of leukocyte stimulating signaling networks during AKI to CKD transition. (A) Umap of the integrated datasets with re-categorized cell type names for Ligand-Receptor analysis. (B) Changes in the standardized interaction scores for Ccl2-Ccr2 ligand-receptor pair between injured proximal tubular cells and interstitial cells. (C) Umap displaying expression levels of Ccl2 and Ccr2 in each group. (D) Heatmap displaying sum of the leukocyte chemotaxis relating L-R interaction scores from each cell types to leukocytes

**Figure 6.**
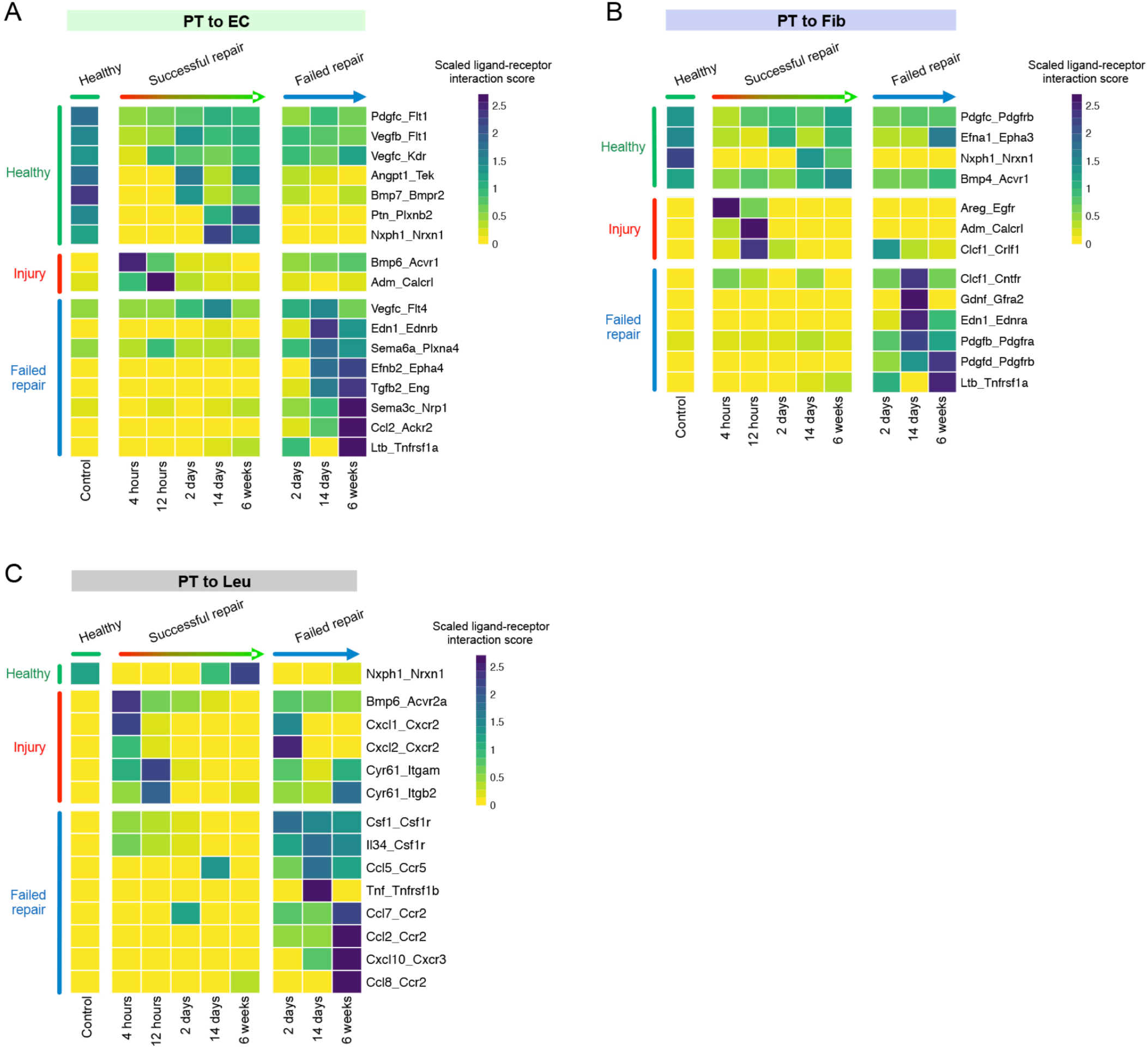
Signaling from proximal tubule to interstitial cell types in health and injury. Heatmaps displaying scaled ligand-receptor interaction scores between proximal tubular cells and interstitial cells (A; PT to endothelial cells, B; PT to fibroblasts, C; PT to leukocytes).

To examine leukocyte chemotactic signaling across cell types more globally, we extracted genes from the “leukocyte chemotaxis” GO term and summed the ligand-interaction score for all cell types across time (Fig. 5D). The strongest scores were seen in endothelium and fibroblasts, with increasing values over time – suggesting ongoing leukocyte signaling even after repair was apparently nearly complete, at six weeks. Consistent with our analysis of Ccl2 signaling, we only observed significant leukocyte chemotactic signaling from epithelia at late timepoints in the injured or FR-PTC clusters (Fig. 5D). These results highlight striking differences in cell types that are promoting inflammation in kidney after injury. In the acute phase, pro-inflammatory fibroblasts and endothelium predominate but in the chronic phase, FR-PTC drive ongoing inflammation.

We then compared proinflammatory and profibrotic signaling from successful repair proximal tubule vs. FR-PTC across time. The pattern was similar whether proximal tubule was signaling to endothelial cells, fibroblasts or leukocytes. For proximal tubule destined to successfully repair, there was very early upregulation of growth factors or cytokines followed by downregulation beginning two days after injury (Fig. 6 A-C). By contrast, the FR-PTC cluster arising at two days after injury upregulated a distinct set of secreted proteins whose expression rose and continued to increase six weeks after injury. Ligands from FR-PTC targeting endothelium include *Edn1*, encoding the potent vasoconstrictor endothelin-1, as well as *Tgfb2* encoding transforming growth factor beta-2 which promotes fibrosis and *Ltb*, encoding lymphotoxin-β which drives inflammatory lymphangiogenesis (31). FR-PTC signaling to fibroblasts included the profibrotic genes *Pdgfrb* and *Pdgfrd* (32). FR-PTC signaling to leukocytes included a variety of pro-inflammatory and pro-fibrotic cytokines including *Csf1, Il34, Ccl5, Tnf, Ccl2, Ccl7, Ccl8* and *Cxcl10*. Consistent with a pro-inflammatory role for FR-PTC, Vcam1+ tubules were surrounded by F4/80+ macrophages at late timepoints after IRI (Fig. S8).

## Discussion

This single nucleus atlas of mouse acute kidney injury will serve as a resource for future studies aimed at understanding cellular responses to kidney injury. Our ability to differentiate between proximal tubule cells that are undergoing successful vs. failed repair allowed the molecular dissection of ligand receptor interactions, signaling pathways and gene regulons that determine whether an injured epithelial cell repairs successfully or not. Whether similar failed repair cell states are shared across organs will be an important question for future studies. Deconvolution of bulk RNA-seq datasets suggests that FR-PTC also exist in human kidney, and increase with age. Whether these pro-inflammatory cells contribute to the well-described age-associated decline in kidney function is another open question. Our results suggest that targeting these pro-inflammatory FR-PTC may reduce chronic inflammation and fibrosis after injury, improving repair.

## Materials and Methods

### Animals

All mouse experiments were performed according to the animal experimental guidelines issued by the Animal Care and Use Committee at Washington University in St. Louis. C57BL/6J (JAX Stock # 000664) were purchased from Jackson Laboratories (Bar Harbor, ME).

### Surgery

For bilateral IRI, 8-10 weeks old male mice were anesthetized with isoflurane and buprenorphine SR was administered for pain control. Body temperature was monitored and maintained at 36.5-37.5°C throughout the procedure. Bilateral flank incisions were made and the kidneys were exposed. Ischemia was induced by clamping the renal pedicle with a non-traumatic microaneurysm clamp (Roboz. Rockville, MD) for 18 minutes. The clamps were subsequently removed and kidneys were returned to the peritoneal cavity. The peritoneal layer was closed with absorbable suture and the flank incisions were closed with wound clips. Control mice underwent sham surgery.

### Mouse kidney samples

Mice were euthanized with isoflurane, blood was collected and the left ventricle was perfused with PBS. For snRNA-seq, kidneys were snap-frozen with liquid nitrogen. For frozen sections, kidneys were fixed with 4% paraformaldehyde for 2 h on ice, incubated in 30% (vol/vol) sucrose at 4 °C overnight, and embedded in optimum cutting temperature compound (Sakura FineTek) to cut 7-µm sections. For paraffin sections, kidneys were fixed with 10% (vol/vol) formalin and paraffin-embedded to cut 4-µm sections. Immunofluorescence protocols and antibodies are detailed below.

### BUN measurement

BUN measurement was done using the QuantiChrom Urea Assay kit as per the manufacturer’s protocol.

### snRNA-seq

Single nuclei isolation from tissue was performed as previously described (33). Briefly, nuclei were isolated with Nuclei EZ Lysis buffer (Sigma #NUC-101) supplemented with protease inhibitor (Roche #5892791001) and RNase inhibitor (Promega #N2615, Life Technologies #AM2696). Samples were cut into <2 mm pieces and homogenized using a Dounce homogenizer (Kimble Chase #885302-0002) in 2ml of ice-cold Nuclei EZ Lysis buffer and incubated on ice for 5 min with an additional 2ml of lysis buffer. The homogenate was filtered through a 40-µm cell strainer (pluriSelect #43-50040-51) and then centrifuged at 500 x for 5 min at 4 °C. The pellet was resuspended and washed with 4 ml of the buffer and incubated on ice for 5 min. After another centrifugation, the pellet was resuspended in Nuclei Suspension Buffer (1x PBS, 1% BSA, 0.1% RNase inhibitor), filtered through a 5-µm cell strainer (pluriSelect 43-50005). Nuclei were counted on hemocytometers (InCYTO C-chip) and partitioned into each droplet with a barcoded gel bead using the 10X Chromium instrument (10X Genomics, Pleasanton, CA). Single nuclei were lysed and RNAs were reverse transcribed into cDNA within each droplet. After breaking the emulsion, cDNAs were amplified and fragmented followed by the addition of Illumina adapters using Single Cell 3’ Library & Gel Bead Kit (v2). Samples were indexed and sequenced on the S4 flow cell of NovaSeq 6000 (Illumina).

### Data processing of snRNA-seq libraries

snRNA-seq data was processed with zUMIs as previously described (34). Briefly, low-quality barcodes and UMIs were filtered out using the internal read filtering algorithm, then mapped to the mouse reference genome (mm10) using STAR 2.5.3a. Next, zUMIs quantified the reads that were uniquely mapped to exonic, intronic or intergenic region of the genome and inferred the true barcodes that mark nuclei by fitting a k-dimensional multivariate normal distribution with mclust package. Finally, a UMI count table utilizing both exonic and intronic reads were generated for downstream analysis. The whole data processing was executed by running the script on the facilities of the Washington University Center for High Performance Computing, which were partially provided through NIH grant S10 OD018091.

### General strategy of snRNA-seq data analysis

Seurat v3 was used for downstream analyses including, normalization, scaling, clustering of nuclei. We analyzed each batch of mouse sample separately and excluded nuclei with less than 150 or more than 8,000 genes detected. We also excluded nuclei with a relatively high percentage of UMIs mapped to mitochondrial genes (>=0.01) and ribosomal genes (>=0.06). Subsequently, we applied SoupX (35) to remove ambient RNA, because different batches can be affected by different levels of ambient RNA. Briefly, ambient RNA expression is estimated from the empty droplet pool (10 UMI or less) with setting “nonExpressedGeneList” to hemoglobin genes followed by removing ambient RNA counts using “adjustCounts” function with default parameters, according to the tutorial from SoupX package (https://cdn.rawgit.com/constantAmateur/SoupX/master/doc/pbmcTutorial.html). In parallel, we used Scrublet (36) to remove doublets. After merging all mouse data, we log-normalized and scaled the data to remove unwanted sources of variation driven by the number of detected UMIs, performed dimension reduction, clustering and sub-clustering. In addition, after clustering or sub-clustering, we performed curated doublet removal based on known lineage specific markers.

### Clustering and dimension reduction

We first combined all mouse datasets. The highly variable genes for principal component analysis were obtained by identifiying the top 500 variable genes from each dataset with FindVariableFeatures () and merging the list, then performed principal component analysis (“RunPCA” function). Combined mouse datasets were integrated using the “RunHarmony” function in the Harmony package. Clustering and UMAP were performed in Seurat using the ‘harmony’ data type as the dimensional reduction type (i.e. reduction.type=‘harmony’). Marker genes were identified from each aligned cell type using the FindAllMarkers function in Seurat. Cluster reassignment was performed based on manual review of lineage-specific marker expression.

### Time course analysis of proximal tubular cells

We subsetted mouse proximal tubular cell clusters, then performed clustering without Harmony integration. The highly variable genes for principal component analysis were obtained by identifying the top 300 variable genes from each datasets with FindVariableFeatures () and merging the list. We then performed principal component analysis (“RunPCA” function), clustering and UMAP.

### Pseudotemporal analysis

Pseudotemporal analysis was performed using Monocle2. We ordered the cells onto a pseudotime trajectory based on the union of highly variable genes in a set of PCs that were previously used for time course analysis of proximal tubular cells. Next, we defined the branch-dependent genes by BEAM function in Monocle2, then cataloged them into 2 clusters in a pseudotime manner. Finally, we performed gene ontology analysis of each gene cluster.

### Gene regulatory network analysis on proximal tubular cells

We used SCENIC for gene regulatory network analysis. In brief, we generated co-expression networks of mouse proximal tubular nuclei data via GRNBoost2. We then utilized the SCENIC package to generate cell regulatory networks from mouse proximal tubular nuclei data, with the mouse mm10 genome for cis-regulatory analysis. We used two gene-motif rankings: 10 kb around the TSS or 500 bp up- and 100 bp downstream the transcriptional start site, which obtained from “https://resources.aertslab.org/cistarget/”.

### Single cell deconvolution

We used BSeq-sc to estimate the proportion of each PT subtype identified from snRNA-seq in the previously reported bulk RNA-seq data as previously described (37). Briefly, the marker genes for each PT subtype and the RPKM normalized gene expression matrix from bulk RNA-seq were used as input according to the tutorial from BSeq-sc package (https://shenorrlab.github.io/bseqsc/vignettes/bseq-sc.html).

### Ligand-receptor interaction analysis

To study ligand-receptor interactions across cell types, we used a draft network (38), and defined an interaction score as previously described with a slight modification (30). In brief, first we eliminated ligand-receptor pairs that include laminin and collagen genes as ligands, then considered only ligands and receptors expressed in more than 0.5% of the nuclei in the specific cell type. Next we defined the interaction score as the product of the average expression of a ligand in a cell type at a time point and the average expression of its cognate receptor of another cell type at same time point. We then standardized each ligand-receptor interaction score by taking the distance between the interaction score and the mean interaction score in units of standard deviations.

### Immunofluorescence

Kidneys were fixed in 4% paraformaldehyde (Electron Microscopy Services), cryoprotected in 30% sucrose solution overnight and embedded in optimum cutting temperature (OCT) compound (Tissue Tek). Kidneys were cryosectioned at 7 micron thickness and mounted on Superfrost slides (Thermo Fisher Scientific). Sections were washed with PBS (3 times, 5 minutes each), then blocked with 10% normal goat serum (Vector Labs), permeabilized with 0.2% Triton X-100 in PBS and then stained with primary antibody specific for Cy3-conjugated anti-αSMA (1:400, Sigma, #C6198), rat anti-PDGFRβ (1:200, eBioscience, #16-1402), rabbit anti-CD31 (1:200, Abcam, #ab28364) and rat anti-F4/80 (1:200, Abcam, #ab6640). Secondary antibodies included AF488-, Cy3-, or Cy5-conjugated (Jackson ImmunoResearch). Then, sections were stained with DAPI (40,60-diamidino-2-phenylindole) and mounted in Prolong Gold (Life Technologies). Images were obtained by confocal microscopy (Nikon C2+ Eclipse; Nikon, Melville, NY)

## Acknowledgments

These experiments were funded by NIH/NIDDK DK103740 and DK103225 and a Chan Zuckerberg Initiative Seed Network Grant.

