## Supplemental Materials for "Cell profiling of mouse acute kidney injury reveals conserved cellular responses to injury"

**This PDF file includes:**

Supplementary text
Figures S1 to S6
Legends for Datasets S1 to S3
SI References

**Other supplementary materials for this manuscript include the following:**

Datasets S1 to S3

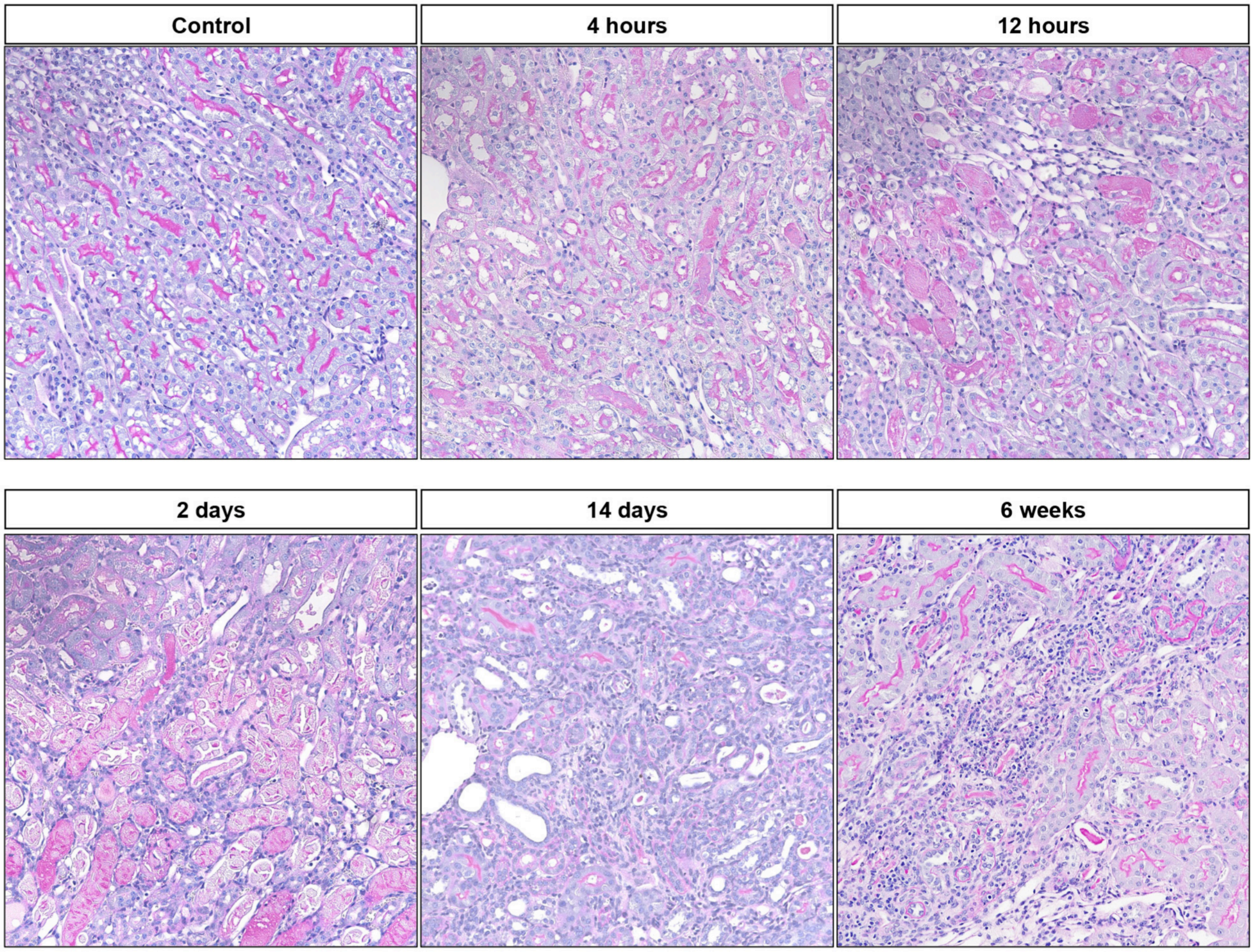


**Fig. S1. Mouse kidney histology.** Representative light microscopy of Periodic acid-Schiff stained mouse kidney over the AKI time course.

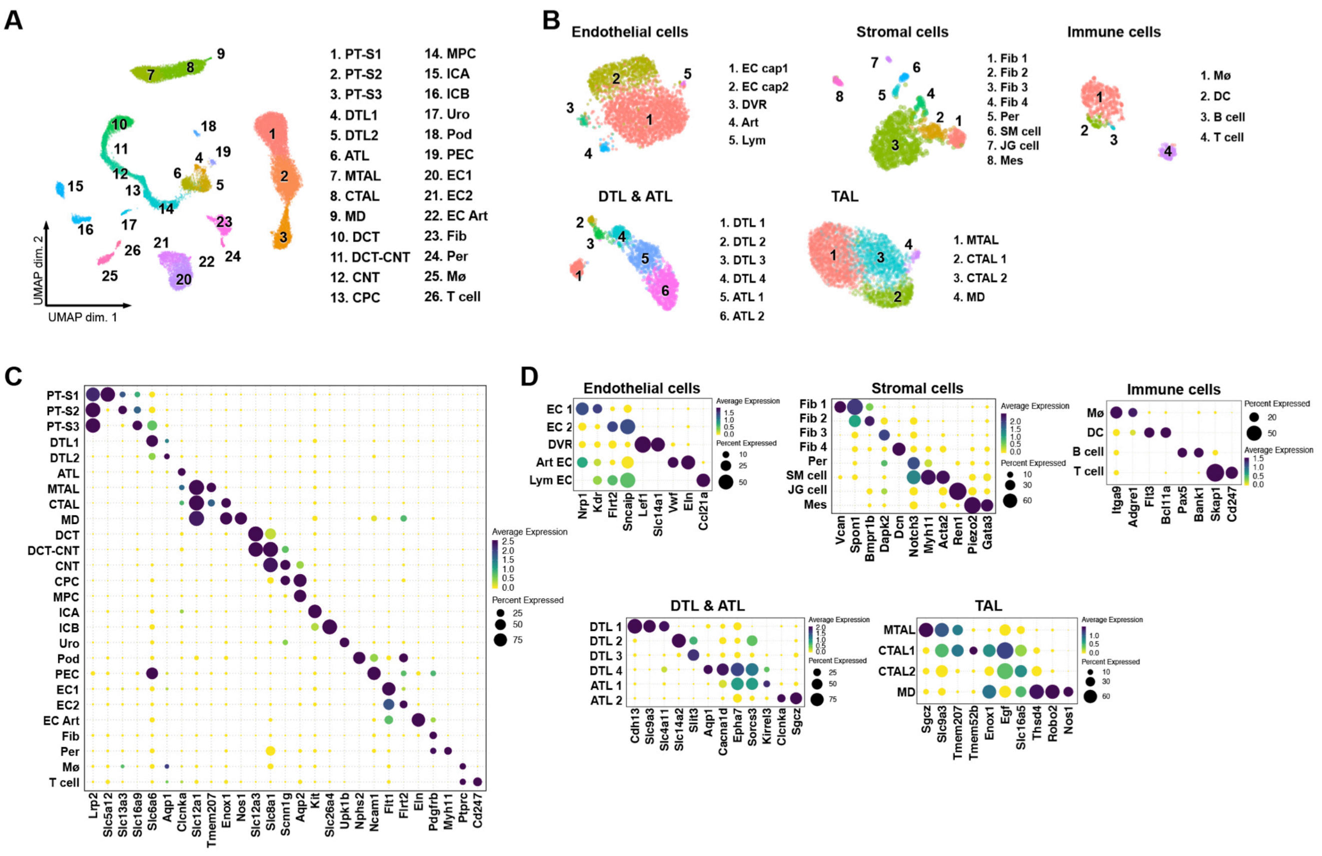


**Fig. S2. Single nucleus RNA sequencing of healthy kidney** (A, B) Umap plots of healthy mouse kidney dataset displaying main- and sub-clusters. PT-S1, S1 segment of proximal tubule; PT-S2, S2 segment of proximal tubule; PT-S3, S3 segment of proximal tubule; DTL, descending limb of loop of henle; ATL, thin ascending limb of loop of henle; MTAL thick ascending limb of loop of henle in medulla; CTAL thick ascending limb of loop of henle in cortex; MD, macula densa; DCT, distal convoluted tubule; CNT, connecting tubule; CPC; principle cells of collecting duct in cortex; MPC; principle cells of collecting duct in medulla; ICA, type A intercalated cells of collecting duct; ICB, type B intercalated cells of collecting duct; Uro, urothelium; Pod, podocytes; PEC, parietal epithelial cells; EC, endothelial cells; Fib fibroblasts; Per, pericytes; Mø, macrophages;  DC, dendritic cells; DVR, descending vasa recta; Art EC, arteriole EC; Lym EC, lymphatic EC; SM, smooth muscle; JG, juxtaglomerular; Mes, mesangial cells. (C, D) Dotplots of heatlhy mouse kidney dataset showing gene expression patterns of cluster-enriched markers.

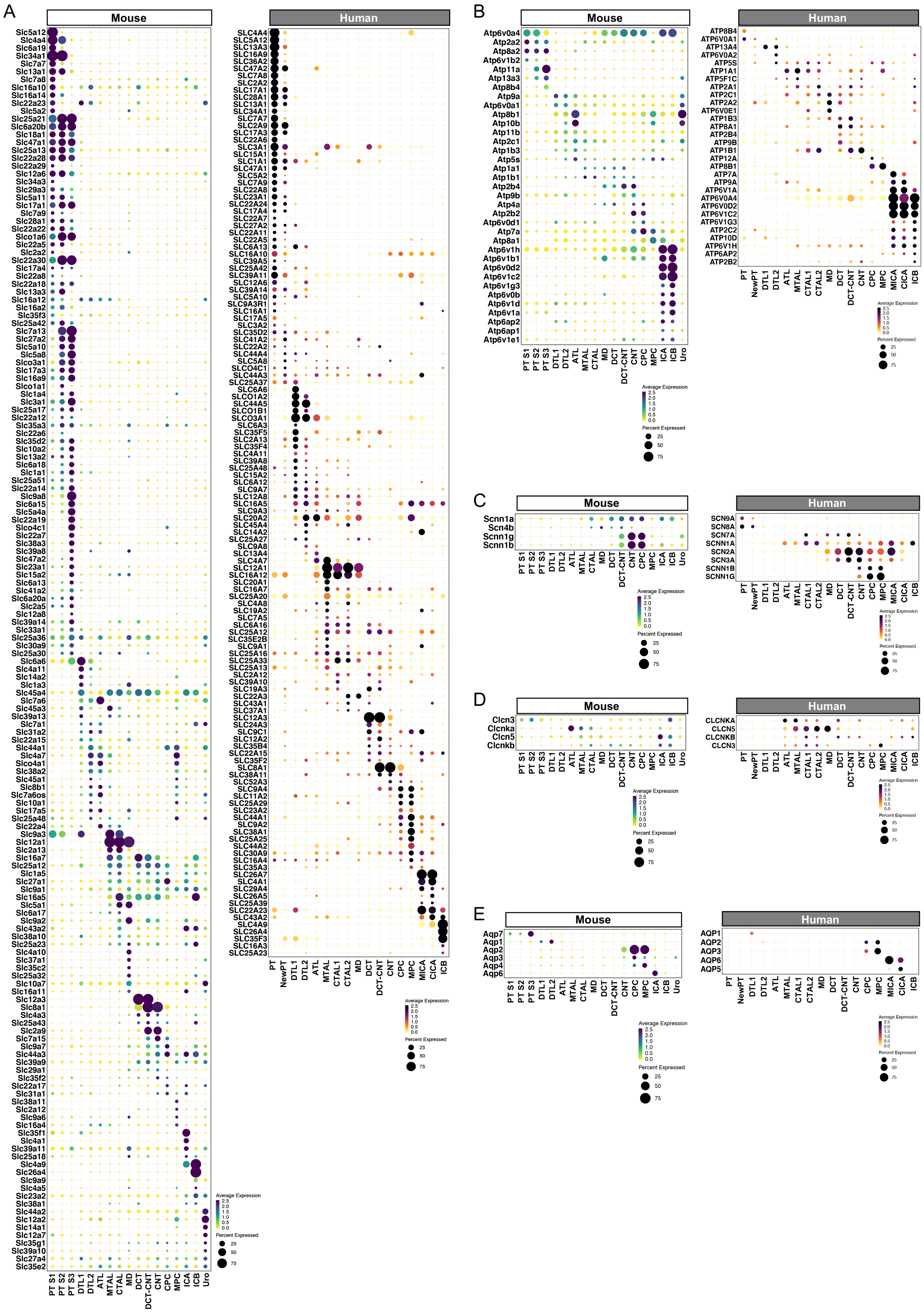

A
Mouse
Human
B
Mouse
Human
Atp6v0a4
Atp2a2
Atp8a2
Atp6v1b2
Atp11a
Atp13a3
Atp8b4
Atp9a
Atp6v0a1
Atp8b1
Atp10b
Atp11b
Atp2c1
Atp1b3
Atp5s
Atp1a1
Atp1b1
Atp2b4
Atp9b
Atp4a
Atp2b2
Atp6v0d1
Atp7a
Atp8a1
Atp6v1h
Atp6v1b1
Atp6v0d2
Atp6v1c2
Atp6v1g3
Atp6v0b
Atp6v1d
Atp6v1a
Atp6ap2
Atp6ap1
Atp6v1e1
ATP8B4
ATP6V0A1
ATP13A4
ATP6V0A2
ATP5S
ATP1A1
ATP5F1C
ATP2A1
ATP2C1
ATP2A2
ATP6V0E1
ATP1B3
ATP8A1
ATP2B4
ATP9B
ATP1B1
ATP12A
ATP8B1
ATP7A
ATP9A
ATP6V1A
ATP6V0A4
ATP6V0D2
ATP6V1C2
ATP6V1G3
ATP2C2
ATP10D
ATP6V1H
ATP6AP2
ATP2B2
C
Mouse
Human
Scnn1a
Scn4b
Scnn1g
Scnn1b
SCN9A
SCN8A
SCN7A
SCNN1A
SCN2A
SCN3A
SCNN1B
SCNN1G
D
Mouse
Human
Clcn3
Clcnka
Clcn5
Clcnkb
CLCNKA
CLCN5
CLCNKB
CLCN3
E
Mouse
Human
Aqp7
Aqp1
Aqp2
Aqp3
Aqp4
Aqp6
AQP1
AQP2
AQP3
AQP6
AQP5
PT S1
PT S2
PT S3
DTL1
DTL2
ATL
MTAL
CTAL
MD
DCT
DCT-CNT
CNT
CPC
MPC
ICA
ICB
Uro
PT
NewPT
CTAL1
CTAL2
MICA
CICA
Average Expression
2.5
2.0
1.5
1.0
0.5
0.0
Percent Expressed
25
50
75

**Fig. S3. Cell-specific expression of transporters, ATPases and channels in mouse kidney.** (A) Expression of all detected solute linked carriers across the nephron in mouse and human kidney. (B) Expression of all detected ATP-driven transporters across the mouse and human nephron. (C) Expression of epithelial sodium channels across mouse and human nephron. (D) Expression of chloride channels across mouse and human nephron. (E) Expression of aquaporins across mouse and human nephron. PT-S1, S1 segment of proximal tubule; PT-S2, S2 segment of proximal tubule; PT-S3, S3 segment of proximal tubule; DTL, descending limb of loop of henle; ATL, thin ascending limb of loop of henle; MTAL thick ascending limb of loop of henle in medulla; CTAL thick ascending limb of loop of henle in cortex; MD, macula densa; DCT, distal convoluted tubule; CNT, connecting tubule; CPC; principle cells of collecting duct in cortex; MPC; principle cells of collecting duct in medulla; ICA, type A intercalated cells of collecting duct; MICA, medullary ICA; CICA, cortical ICA ; ICB, type B intercalated cells of collecting duct; Uro, urothelium.

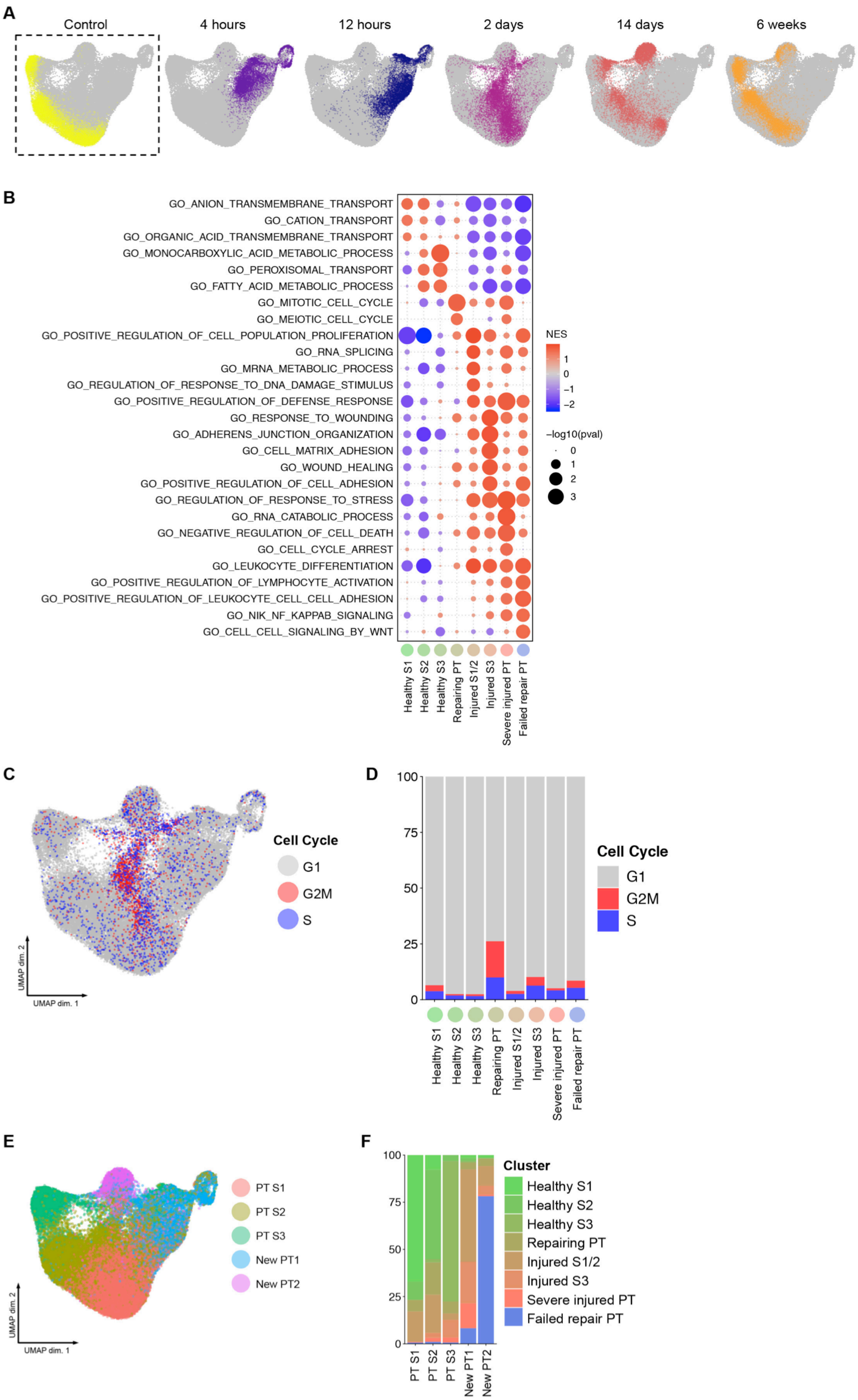


**Fig. S4.** Time-course analysis of proximal tubular cells. (A) Projection of timepoints after IRI for all proximal tubule cells. (B) Gene ontology enrichment analysis of proximal tubule subclusters. (C) Projection of cell cycle states onto Umap space. (D) Fraction of proximal tubule cells in the

G2M, S or G1 cell cycle across subclusters. (E) Projection of main-clustering onto UMAP of all proximal tubule cells. (F) Fraction of proximal tubule cells in the subclusters across main-clusters.

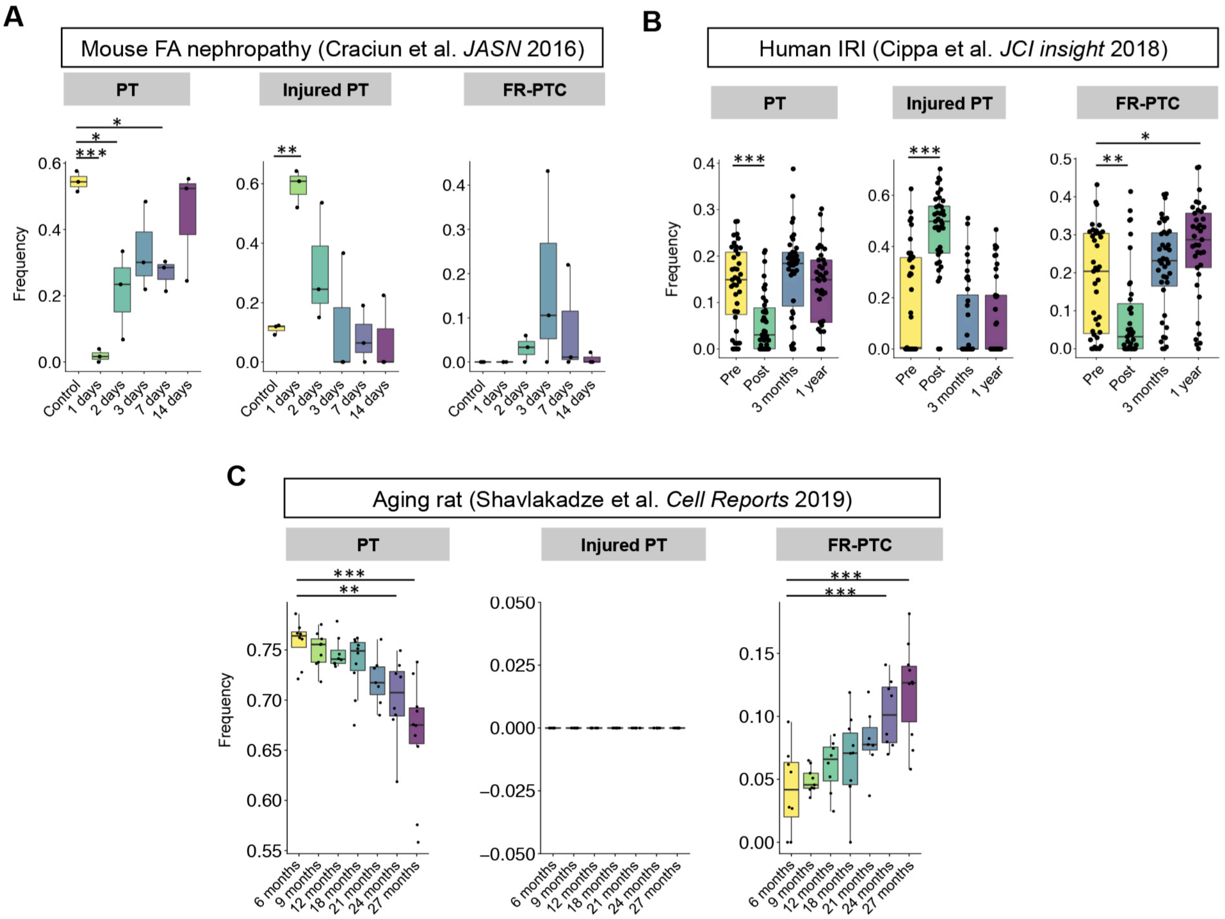


**Fig. S5. Deconvolution of PT subtype composition in previous bulk RNA-seq datasets.** (A) Mouse folic acid nephropathy, (B) Human IRI, (D) Aging rat. **P* < 0.05; ***P* < 0.01; ****P* < 0.001, one-way ANOVA with post hoc Dunnett's multiple comparisons test.

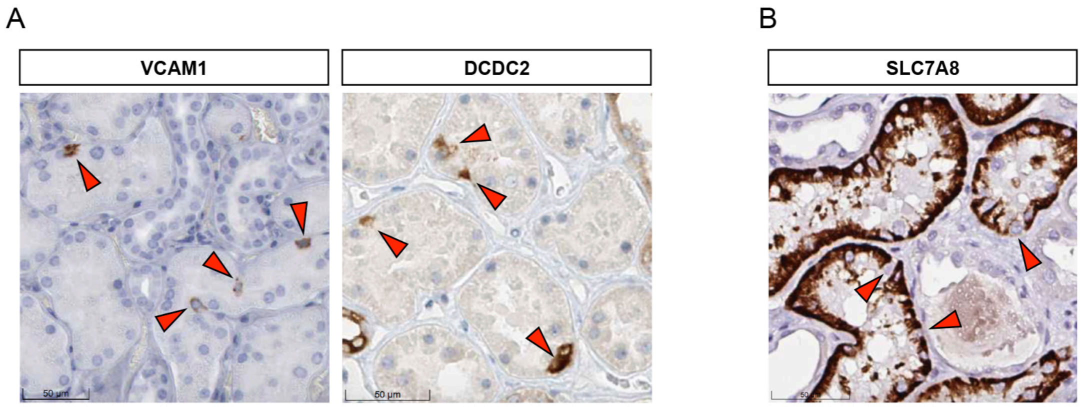


**Fig. S6.** Immunohistochemical profile of individual scattered proximal tubule cells in the Human Protein Atlas. (A) Scattered cells show increased expression of VCAM1 and DCDC2. (B) Scattered cells show decreased expression of SLC7A8.

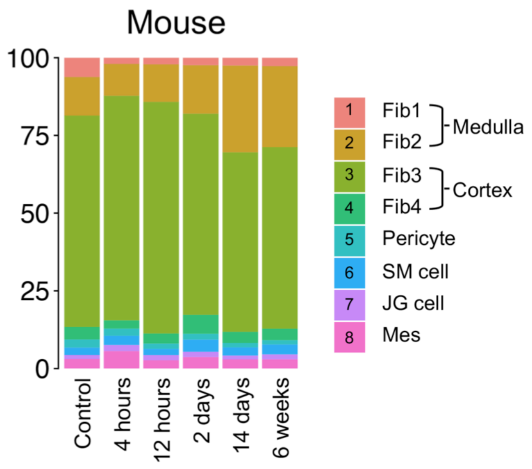


**Fig. S7.** Medullary fibroblasts increase after IRI. Bar plot displaying composition of groups by clusters in stromal cells.

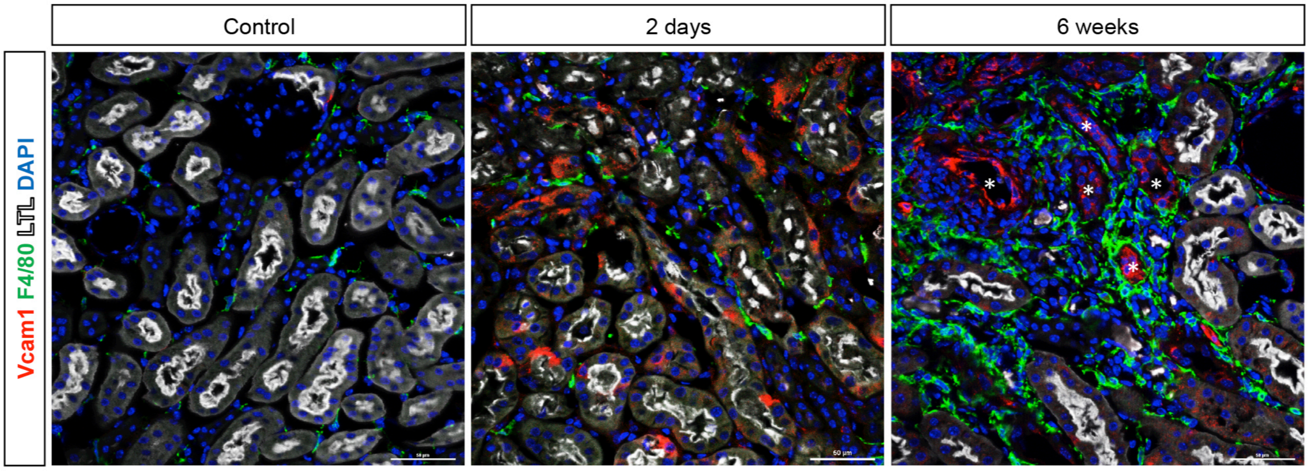


**Fig. S8.** Representative images of immunofluorescence staining for VCAM1 (red), F4/80 (green) and LTL (white). * Vcam1-positive tubules surrounded by F4/80-positive interstitial macrophages.

**Dataset S1 (separate file).** Differential expression of cell-type-specific genes in healthy mouse kidney.

**Dataset S2 (separate file).** Differential expression of proximal tubule-specific genes between healthy control and injured/dedifferentiated cells with accompanying GO ontology enrichment analysis.

**Dataset S3 (separate file).** Regulon activity in the healthy control and injured/dedifferentiated proximal tubule determined by SCENIC.
